# A deep neural network estimation of brain age is sensitive to cognitive impairment and decline

**DOI:** 10.1101/2023.08.10.552494

**Authors:** Yisu Yang, Aditi Sathe, Kurt Schilling, Niranjana Shashikumar, Elizabeth Moore, Logan Dumitrescu, Kimberly R. Pechman, Bennett A. Landman, Katherine A. Gifford, Timothy J. Hohman, Angela L. Jefferson, Derek B. Archer

## Abstract

The greatest known risk factor for Alzheimer’s disease (AD) is age. While both normal aging and AD pathology involve structural changes in the brain, their trajectories of atrophy are not the same. Recent developments in artificial intelligence have encouraged studies to leverage neuroimaging-derived measures and deep learning approaches to predict brain age, which has shown promise as a sensitive biomarker in diagnosing and monitoring AD. However, prior efforts primarily involved structural magnetic resonance imaging and conventional diffusion MRI (dMRI) metrics without accounting for partial volume effects. To address this issue, we post-processed our dMRI scans with an advanced free-water (FW) correction technique to compute distinct FW-corrected fractional anisotropy (FA_FWcorr_) and FW maps that allow for the separation of tissue from fluid in a scan. We built 3 densely connected neural networks from FW-corrected dMRI, T1-weighted MRI, and combined FW+T1 features, respectively, to predict brain age. We then investigated the relationship of actual age and predicted brain ages with cognition. We found that all models accurately predicted actual age in cognitively unimpaired (CU) controls (FW: r=0.66, *p*=1.62×10^−32^; T1: r=0.61, *p*=1.45×10^−26^, FW+T1: r=0.77, *p*=6.48×10^−50^) and distinguished between CU and mild cognitive impairment participants (FW: *p*=0.006; T1: *p*=0.048; FW+T1: *p*=0.003), with FW+T1-derived age showing best performance. Additionally, all predicted brain age models were significantly associated with cross-sectional cognition (memory, FW: β=−1.094, *p*=6.32×10^−7^; T1: β=−1.331, *p*=6.52×10^−7^; FW+T1: β=−1.476, *p*=2.53×10^−10^; executive function, FW: β=−1.276, *p*=1.46×10^−9^; T1: β=−1.337, *p*=2.52×10^−7^; FW+T1: β=−1.850, *p*=3.85×10^−17^) and longitudinal cognition (memory, FW: β=−0.091, *p*=4.62×10^−11^; T1: β=−0.097, *p*=1.40×10^−8^; FW+T1: β=−0.101, *p*=1.35×10^−11^; executive function, FW: β=−0.125, *p*=1.20×10^−10^; T1: β=−0.163, *p*=4.25×10^−12^; FW+T1: β=−0.158, *p*=1.65×10^−14^). Our findings provide evidence that both T1-weighted MRI and dMRI measures improve brain age prediction and support predicted brain age as a sensitive biomarker of cognition and cognitive decline.

## 1. Introduction

Alzheimer’s disease (AD) is a progressive neurodegenerative disorder whose greatest known risk factor is advancing age. Both normal aging and AD are accompanied by structural changes in the brain, but they follow distinct trajectories. Specifically, healthy aging typically exhibits global reductions in gray matter volume^1,2^ characterized by volume loss in frontal and temporal lobes^3,4^ and enlargement of ventricles^3,5^, whereas AD-related brain atrophy typically starts in the hippocampus and gradually spreads to the entire brain^6,7^. Additionally, studies have shown that AD brains undergo deterioration more rapidly than healthy brains^8^. Given these differences, there arose recent efforts of using neuroimaging-derived measures of gray matter volume from T1-weighted magnetic resonance imaging (MRI) and white matter microstructure from diffusion MRI (dMRI) to predict an individual’s “brain age” via machine learning approaches^9–12^, which can differ from their chronological age and predict cognitive decline^13–15^. These models were trained on cognitive unimpaired individuals to learn common patterns in healthy aging, which then allowed them to detect aging-related abnormalities such as those associated with AD. A larger difference between brain age and chronological age indicates that the individual is on an accelerated trajectory compared with normal aging and is typically seen in individuals with cognitive impairment (e.g., mild cognitive impairment [MCI], AD)^16–18^, suggesting the potential of brain age as a sensitive biomarker along the AD continuum. Moreover, the development of the free-water (FW) correction post-processing technique^19^ has enabled the partition of a conventional fractional anisotropy (FA) map into a FW-corrected FA map (FA_FWcorr_) and a FW map; the FA_FWcorr_ and FW metrics individually describe tissue and fluid, thereby enhancing the biological specificity of dMRI scans. Recently, our group has demonstrated that abnormal FW-corrected dMRI metrics are associated with higher rates of longitudinal cognitive decline and diagnosis along the AD clinical continuum^20,21^. These findings suggest that incorporating FW-corrected metrics into models of predicted brain age may provide more sensitive associations with cognitive impairment and decline.

The present study leveraged neuroimaging data from a longitudinal cohort of aging to build three densely connected neural networks using FW-corrected dMRI, T1-weighted MRI, and combined FW+T1 features to predict participant brain age. To evaluate model performance, we examined the relationship between predicted brain age and chronological age. We then investigated the association between predicted brain age and two domains of cognition (memory and executive function performance at baseline and over time). We hypothesized that FW-, T1-, and FW+T1-derived models would all accurately predict participant brain age, with the FW+T1-derived model showing the best performance as it incorporates both gray and white matter regions. We also hypothesized that all predicted brain age models would predict baseline and longitudinal memory and executive function performance, with FW+T1-derived brain age showing the strongest associations.

## 2. Methods

### 2.1. Participants

All data leveraged in the present study were obtained from the Vanderbilt Memory and Aging Project (VMAP)^22^, a longitudinal observational study that was launched in 2012 and recruited individuals 60 years and older who speak English, have adequate auditory and visual capacity for testing, and have a stable study partner. Participants underwent comprehensive neuropsychological assessment and were categorized into cognitively unimpaired (CU) or MCI status; MCI participants were age-, sex-, and race-matched with CU controls. Cognitive (memory, executive function) measures were obtained from all participants and neuroimaging (T1-weighted MRI, dMRI) measures were obtained from a subset of participants. Only participants who had all necessary cognitive and neuroimaging data were included in the present study (n=295). All protocols for VMAP were approved by the IRB at Vanderbilt University Medical Center and all participants gave voluntary informed consent prior to enrollment. Data from the VMAP cohort can be freely accessed following approval (vmacdata.org). **Table 1** summarizes demographic and clinical information for the present cohort.

**Table 1.** Vanderbilt Memory and Aging Project Cohort Information.

| Measure | Diagnosis at Baseline |  | p-value |
| --- | --- | --- | --- |
|  | CU | MCI |  |
| Cohort Characteristics |  |  |  |
| Number of participants | 168 | 127 | - |
| Total number of visits | 568 | 372 | - |
| Longitudinal follow-up (years) | 3.10 (1.44) | 2.96 (1.43) | 0.230 |
| Demographics and Health Characteristics |  |  |  |
| Age at baseline (years) | 73.10 (7.16) | 73.67 (7.41) | 0.504 |
| Sex (% female) | 42.26 | 42.52 | 1.000 |
| Education (years) | 16.39 (2.48) | 15.09 (2.75) | **<0.001** |
| Race (% non-Hispanic White) | 86.90 | 86.61 | 1.000 |
| APOE-ε4 (% positive) | 29.76 | 44.09 | **0.016** |

Mean (standard error) are provided unless otherwise indicated. Abbreviations: CU, cognitively unimpaired; MCI, mild cognitive impairment; *APOE*, apolipoprotein. Boldface signifies $p < 0.05$ unless otherwise indicated.

### 2.2. Neuroimaging data acquisition and preprocessing

T1-weighted MRI images (repetition time: 8.9 ms, echo time: 4.6 ms, resolution: 1 mm isotropic) were obtained from each participant on 3T Philips Achieva using an 8-channel SENSE reception coil and underwent multi-atlas segmentation to calculate the volumes of 132 regions of interest (ROI)^23^. All measures were normalized by total intracranial volume, calculated as the volumetric sum of all 132 segmented ROIs. dMRI images (resolution: 2 mm isotropic, b-values: 0, 1000 s/ mm^2^, number of directions: 32) were obtained from each participant using the previously described scanner and preprocessed using *PreQual*^24^. FW and FW-corrected metrics were calculated in MATLAB from the preprocessed images, as previously described^19^. The FW and FA_FWcorr_ maps were transformed by a non-linear warp using the ANTs package to create a standardized space representation. Finally, publicly available tractography templates (https://github.com/VUMC-VMAC/Tractography_Templates) were applied to the FW and FA_FWcorr_ maps to quantify white matter microstructure within 48 tracts.

T1-weighted MRI and FW-corrected dMRI metrics (FA_FWcorr_, FW) were harmonized separately using *Longitudinal Combat*^25^ in R (version 4.1.2), controlling for age at baseline, education, sex, race/ethnicity, *APOE*-ε4 positivity, *APOE*-ε2 positivity, and the interaction of age at baseline with time interval from baseline. We also included the random effects of intercept and time interval from baseline for each participant and a batch variable that accounted for all combinations of image acquisition. The batch variable was *scanner x software x coil* for T1 metrics and *site x scanner x protocol* for FW-corrected metrics.

### 2.3. Neuropsychological metrics calculation

Participants completed comprehensive neuropsychological testing administered by experienced technicians which assessed multiple cognitive domains, including memory and executive function. Psychometrically sound memory and executive function composite scores were calculated from item-level data. Longitudinal cognitive measures (memory slope, executive function slope) for each participant were obtained by calculating the random effect coefficient using a linear mixed-effects model where the fixed effect was time interval from baseline and the outcome was composite score.

### 2.4. Brain age prediction model architecture

In the present study, we used a densely connected neural network to predict participants’ brain age based on neuroimaging regions (i.e., features) and created three separate models using FW, T1, and combined FW+T1 features. **Figure 1** shows an overview of model workflow. Each model consists of four layers: an input layer whose dimensions correspond to the number of features (FW: 96 features, T1: 132 features, FW+T1: 228 features), two densely connected layers with rectified linear unit (ReLU) activation whose number of nodes equals half and a quarter of the number of features, respectively, and an output layer with a single node and linear activation for brain age prediction.

**Figure 1.**
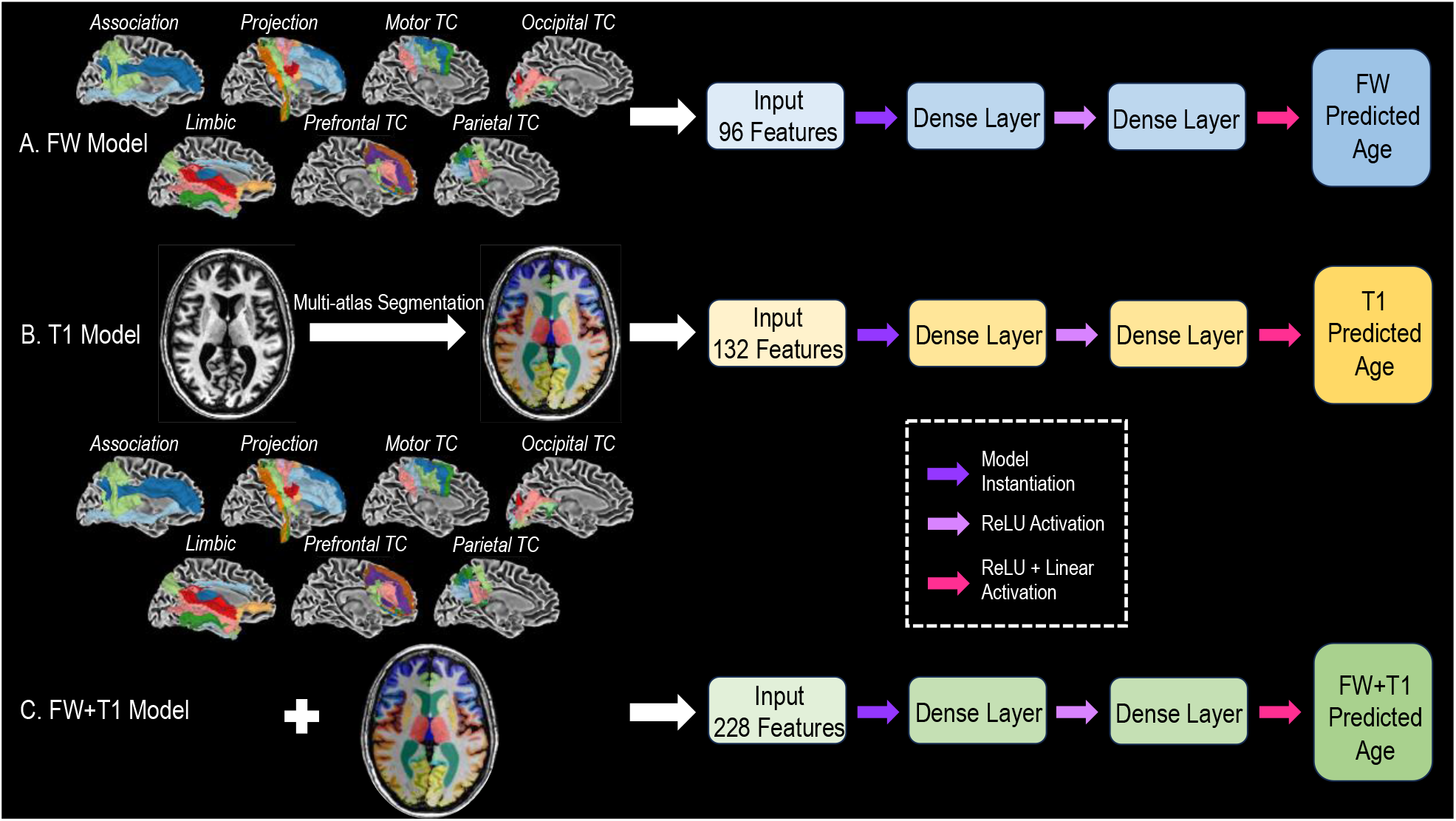
Model workflow for brain age prediction. We created three separate, densely connected neural networks to predict brain age, including FW-derived (**A**), T1-derived (**B**), and FW+T1-derived (**C**) models.

All models were trained on baseline neuroimaging data from the VMAP cohort by subsetting all imaging sessions to the first visit of CU participants. We minimized the loss function as characterized by mean absolute error (MAE) while monitoring the mean squared error (MSE) and root mean squared error (RMSE). We conducted ten-fold cross-validation where 90% of the data were used for training and 10% of the data were reserved for testing in each fold, repeating this process ten times until the entire dataset had been tested only once. Within the training data for each fold, 80% were used to train the model and 20% were used to validate model performance. During each fold, training was stopped when the loss function on the validation dataset had not improved for 15 epochs and only the best model was saved. For each set of features (FW, T1, FW+T1), saved models were compared across folds and the one which yielded the lowest validation loss was selected as the final model. All models were developed in Python (version 3.9.13) using the Keras library (version 2.9.0) with Tensorflow backend (version 2.9.1). We used the three final models to generate FW, T1, and FW+T1 predicted brain ages for all participants (CU, MCI) at all timepoints (baseline, longitudinal follow-ups).

For each model, we computed SHAP (SHapley Additive exPlanation) values for all relevant neuroimaging features to quantify their contribution to age prediction. **Figure 2** shows the top 10 most important features for each model based on mean SHAP value.

**Figure 2.**
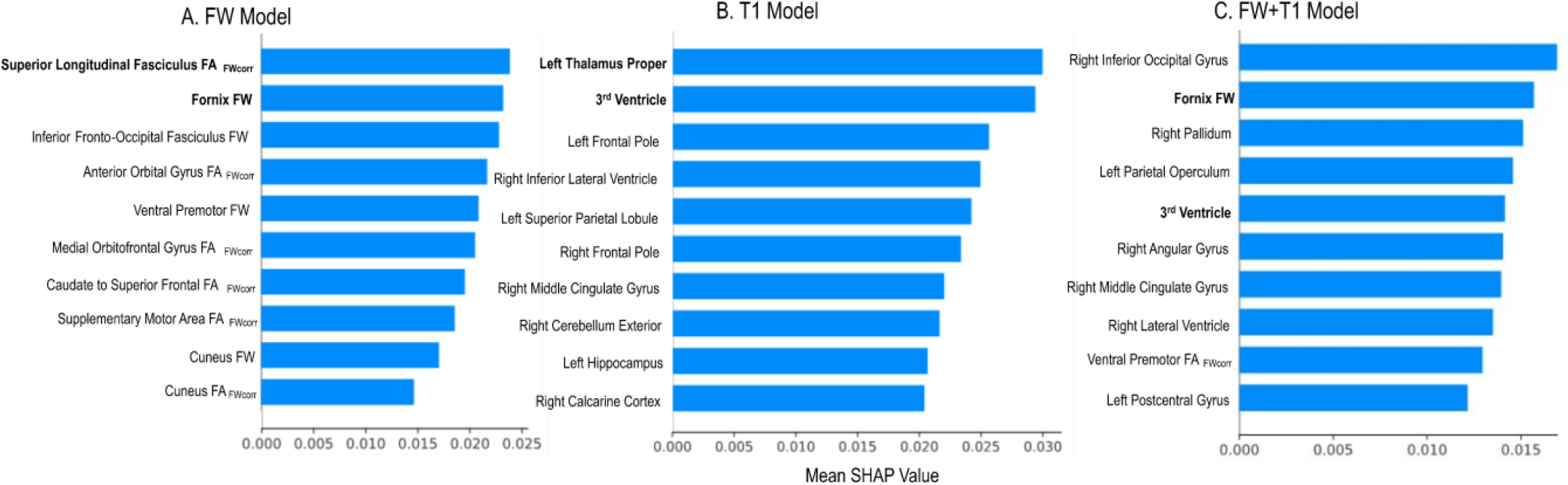
Top 10 most important features for FW-derived (**A**), T1-derived (**B**), and FW+T1-derived (**C**) models. Boldface signifies top features involved in aging and AD, including superior longitudinal fasciculus (SLF) FA_FWcorr_, fornix FW for the FW-derived model and left thalamus proper, 3^rd^ ventricle for the T1-derived model.

### 2.5. Statistical analyses

All statistical analyses were conducted in Python (version 3.9.13) and R (version 4.1.2). We first performed simple linear regression between actual age and each predicted age to assess model performance as well as independent groups t-tests to compare the mean actual age and mean predicted brain ages of CU and MCI participants. We also conducted logistic regression analyses, using actual and each predicted brain age as direct predictor of diagnostic category, then evaluated model performance using area under the receiver operator characteristic curve (ROC-AUC) and DeLong’s test. Next, we conducted a series of linear models and competitive model analyses to assess actual and predicted brain age association with cognition. All models covaried for diagnosis, race/ethnicity, sex, education, and *APOE*-ε4 positivity. Significance was set *a priori* at α=0.05. For baseline cognition, actual age and predicted brain ages (FW, T1, FW+T1) were included in a general linear model individually to determine their main effects on baseline memory and executive function. We then introduced age-by-diagnosis interaction terms to the linear models to investigate the potential modifying effect of age on baseline memory and executive function scores. Finally, we conducted post-hoc competitive model analysis to determine the unique variance in baseline memory and executive function contributed by FW, T1, and FW+T1 predicted brain age, beyond that contributed by covariates and actual age. The described analyses were repeated for longitudinal cognition (longitudinal memory slope, longitudinal executive function slope).

## 3. Results

Participant characteristics of the VMAP cohort are shown in **Table 1**. There were no significant differences in longitudinal follow-up interval, age at baseline, sex, or race between diagnostic groups (CU, MC). The CU group had more years of education and lower *APOE*-ε4 positivity than the MCI group.

### 3.1. Combined model using free-water (FW) and T1 features showed best performance

Figure 3. shows the agreement between predicted brain age measures (FW, T1, FW+T1) and actual age; model performance was characterized using average mean absolute error (MAE_avg_) and average mean squared error (RMSE_avg_) across folds and Pearson’s correlation through ten-fold cross validation. While all predicted brain ages significantly predicted actual age (FW: MAE_avg_=0.115, RMSE_avg_=0.129, r=0.66, *p*=1.62×10^−32^; T1: MAE_avg_=0.106, RMSE_avg_=0.114, r=0.61, *p*=1.45×10^−26^), the combined FW+T1 model yielded the best performance with highest r as well as lowest MAE_avg_ and RMSE_avg_ (MAE_avg_=0.072, RMSE_avg_=0.087, r=0.77, *p*=6.48×10^−50^).

We then compared means of actual age and predicted brain ages between CU and MCI participants. While there was no difference in actual age between CU and MCI groups (age_CU_=73.07±7.24, age_MCI_=72.83±6.92, *p*=0.792), all predicted brain ages for the MCI group were significantly higher than those for the CU group (FW: age_CU_=72.08±5.55, age_MCI_=74.18±6.16, *p*=0.006; T1: age_CU_=67.52±4.96, age_MCI_=68.82±5.27, *p*=0.048), with the combined FW+T1 model showing the largest difference (age_CU_=71.74±5.58, age_MCI_=73.93±5.67, *p*=0.003).

**Figure 3.**
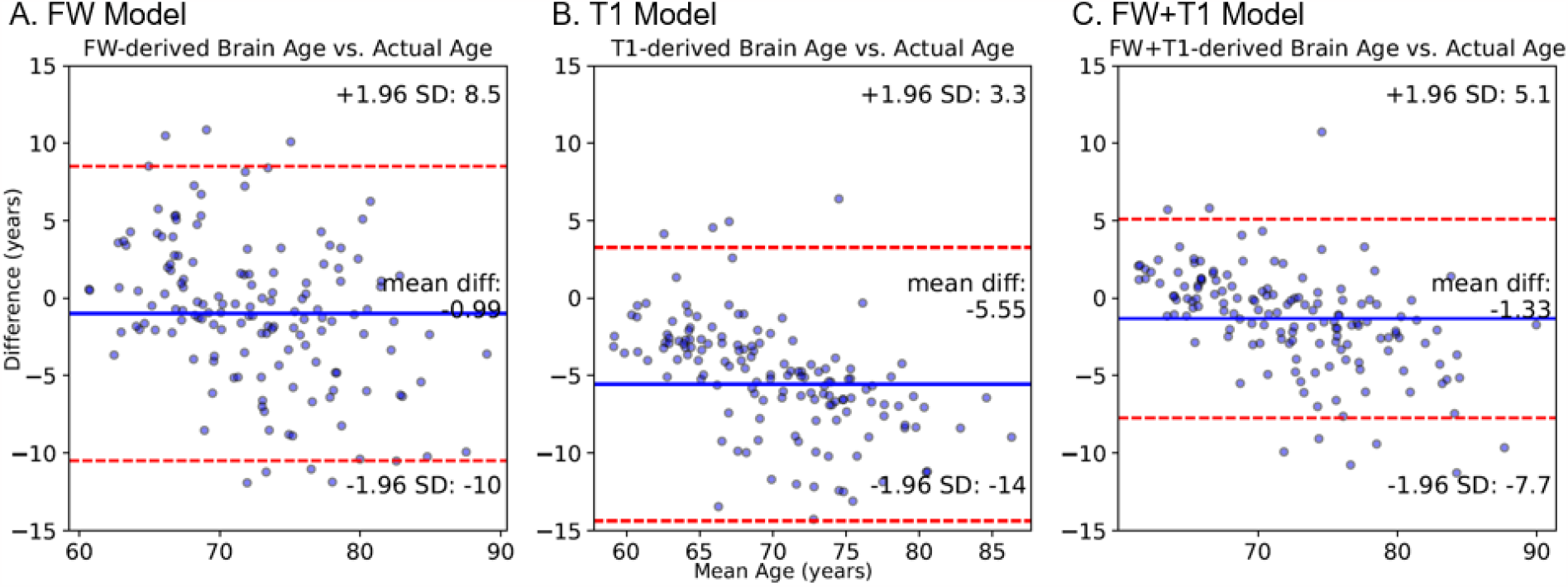
Bland Altman plots for FW-derived age (**A**), T1-derived age (**B**), and FW+T1-derived age (**C**). All models accurately predict age. FW+T1-derived age is most significantly associated with actual age, in comparison with FW-derived age and T1-derived age.

Figure 4. shows the Receiver Operating Characteristic curves for actual and predicted brain ages in predicting diagnostic category (CU, MCI). Pairwise comparisons revealed that ROC-AUC values for all predicted brain ages were significantly greater than that of actual age (FW-actual: *p*=0.003; T1-actual: *p*=0.030; FW+T1-actual: *p*=0.004); however, no differences were found between the predicted brain ages (all *p*>0.05).

### 3.2. Predicted brain age association with baseline cognition

Actual age and predicted brain age (FW-derived, T1-derived, FW+T1-derived) associations with cross-sectional cognition (memory, executive function) are shown in **Figure 5**. While all models significantly predicted memory score at baseline (Actual: R_adj_^2^=0.497, *p*=1.23×10^−34^; FW: R_adj_^2^=0.481, *p*=4.14×10^−33^; T1: R_adj_^2^=0.481, *p*=4.26×10^−33^), the combined FW+T1 model showed the most robust performance (R_adj_^2^=0.513, *p*=2.31×10^−36^). Similarly, all models significantly predicted executive function score at baseline (Actual: R_adj_^2^=0.472, *p*=3.22×10^−32^; FW: R_adj_^2^=0.445, *p*=1.24×10^−29^; T1: R_adj_^2^=0.422, *p*=1.69×10^−27^) and the combined FW+T1 model was the most robust (R_adj_^2^=0.519, *p*=5.81×10^−37^). When examining main effect associations of each respective age variable, we saw that actual age and all predicted brain ages each had a significant main effect on baseline memory score (**Figure 5A**; Actual: β=−1.162, *p*=1.58×10^−8^; FW: β=−1.094, *p*=6.32×10^−7^; T1: β=−1.331, *p*=6.52×10^−7^), with the combined FW+T1 predicted brain age showing the strongest relationship (β=−1.476, *p*=2.53×10^−10^). Likewise, we saw significant age effects for actual and all predicted ages on baseline executive function score (**Figure 5B**; Actual: β=−1.371, *p*=2.98×10^−12^; FW: β=−1.276, *p*=1.46×10^−9^; T1: β=−1.337, *p*=2.52×10^−7^), with the combined FW+T1 predicted brain age showing the strongest relationship (β=−1.850, *p*=3.85×10^−17^). We found no significant interactions between actual or predicted brain ages and diagnostic status on baseline memory or executive function.

**Figure 4.**
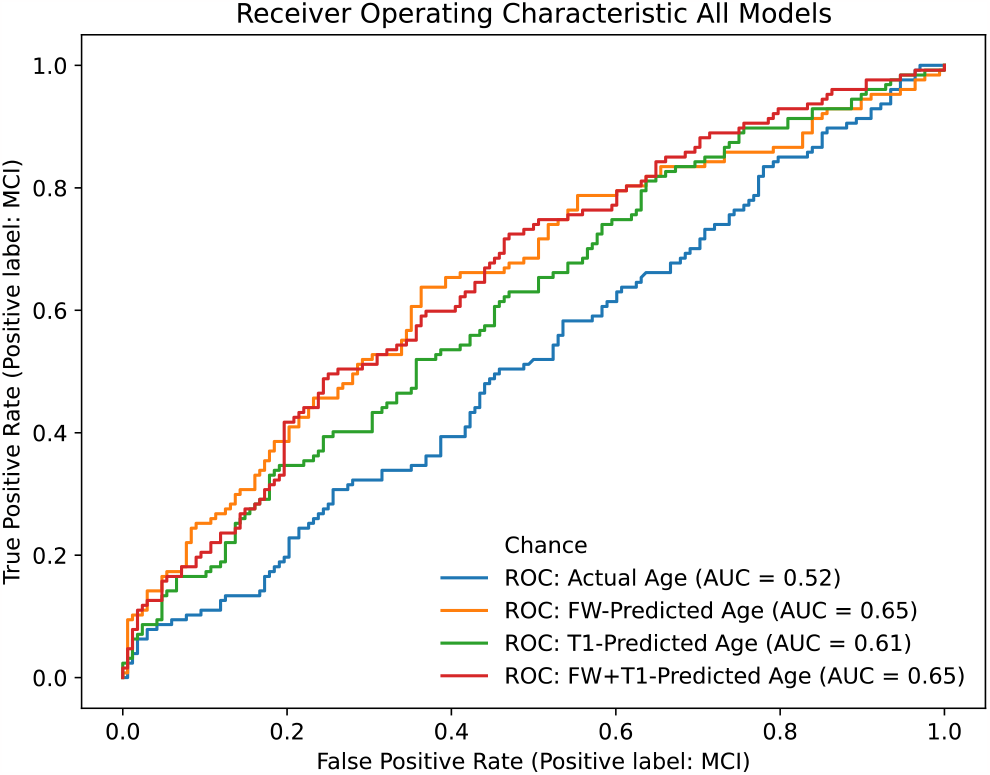
Receiver Operating Characteristic curves for actual, FW-predicted, T1-predicted, and FW+T1-predicted age in predicting diagnostic category. All predicted ages performed significantly better than actual age, but no difference in performance was found between predicted ages.

**Figure 5.**
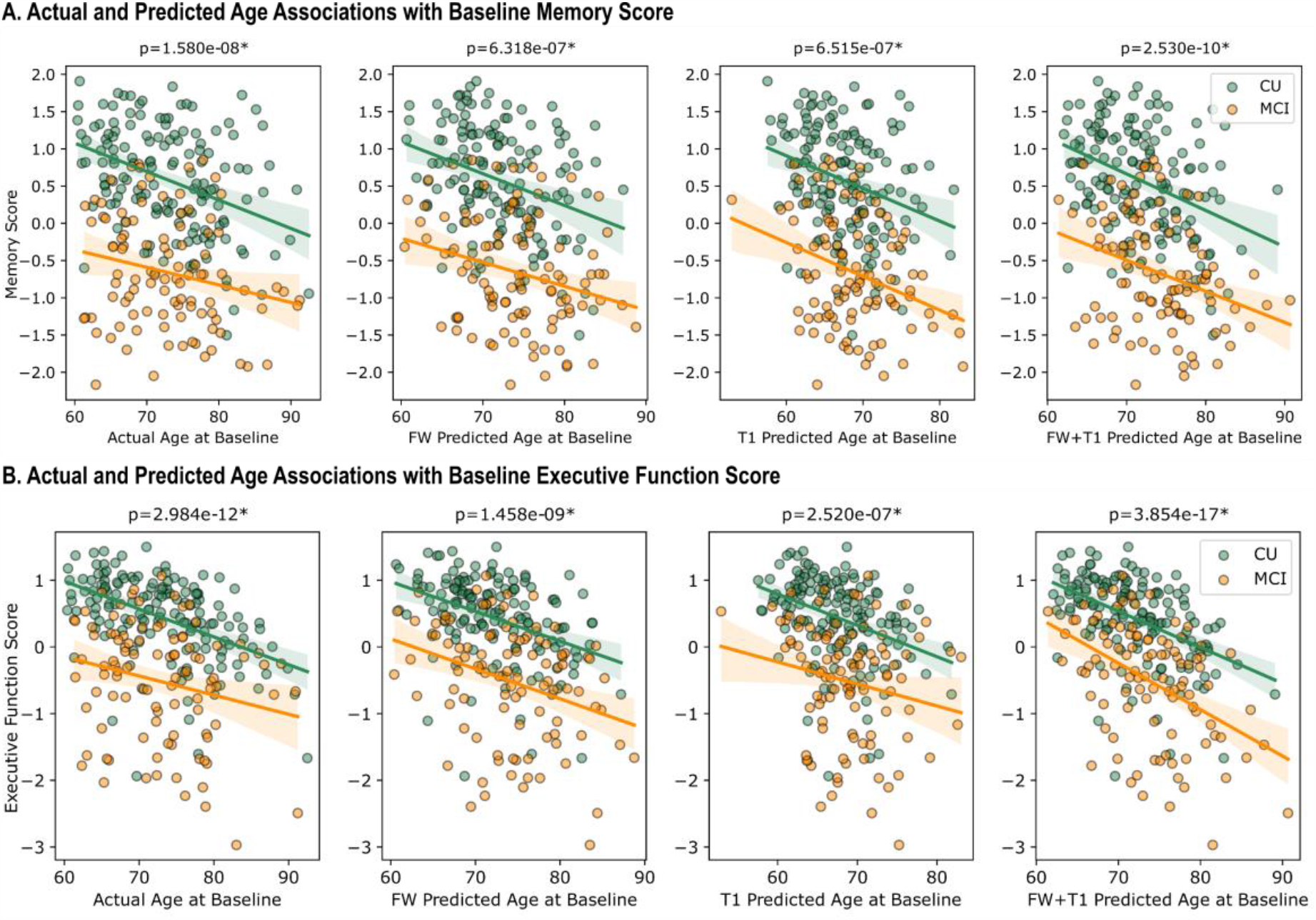
Actual and predicted age associations with baseline cognition. Actual and all derived ages are significantly associated with baseline memory (**A**) and executive function performance (**B**); FW+T1-derived age shows highest associations. Datapoint colors: green=CU; orange=MCI.

**Table 2** summarizes results of the competitive model analysis on cross-sectional cognition. We found that covariates alone explained approximately 43% of the variance in baseline memory score (R_adj_^2^=42.60%) and the addition of actual age led to an increase in overall model performance (ΔR^2^_adj_=6.92%). We then iteratively added predicted brain ages to this model to determine whether FW, T1, or FW+T1 predicted brain age contributed to any unique variance beyond covariates and actual age. While FW and T1 predicted brain ages were not found to be a significant contributor to baseline memory score, we observed that the combined FW+T1 predicted brain age significantly added to the model and led to increased R_adj_^2^ (FW+T1: ΔR_adj_^2^ =1.74%). Similarly, covariates alone explained approximately 36% of the variance in baseline executive function score (R_adj_^2^=35.70%) and the addition of actual age led to a drastic increase in model performance (ΔR^2^_adj_=11.58%). When iteratively adding each predicted brain age to the model to determine its unique contribution beyond covariates and actual age, we observed that both FW and FW+T1 predicted brain age explained additional variance in baseline executive function score, with FW predicted brain age leading to a small increase in R_adj_^2^ (ΔR_adj_^2^ =0.63%) and FW+T1 predicted brain age leading to a large increase in R_adj_^2^ (ΔR_adj_^2^ =4.59%). However, T1 predicted brain age did not provide a significant increase to the model.

**Table 2.** Comparison of Actual and Predicted Age Main Effects on Baseline Cognition.

|  | Baseline Memory Score |  |  |  |  | Baseline Executive Function Score |  |  |  |  |
| --- | --- | --- | --- | --- | --- | --- | --- | --- | --- | --- |
| | $\beta$ | SE | t | p | $\Delta R_{adj}^2$ | $\beta$ | SE | t | p | $\Delta R_{adj}^2$ |
| Covariates + Actual age | -1.162 | 0.199 | -5.852 | **<0.001** | 6.915 | -1.371 | 0.186 | -7.355 | **<0.001** | 11.576 |
| Covariates + actual age + Predicted age |  |  |  |  |  |  |  |  |  |  |
| FW | -0.474 | 0.287 | -1.652 | 0.100 | 0.358 | -0.531 | 0.269 | -1.978 | **0.049** | 0.630 |
| T1 | -0.643 | 0.327 | -1.964 | 0.051 | 0.590 | -0.376 | 0.308 | -1.219 | 0.224 | 0.106 |
| FW+T1 | -1.132 | 0.365 | -3.105 | **0.002** | 1.743 | -1.628 | 0.333 | -4.892 | **<0.001** | 4.585 |

### 3.3. Predicted brain age association with longitudinal cognition

Actual age and predicted brain age associations with longitudinal cognition are shown in **Figure 6**. While all models significantly predicted longitudinal memory slope (Actual: R_adj_^2^=0.427, *p*=5.08×10^−28^; FW: R_adj_^2^=0.439, *p*=4.90×10^−29^; T1: R_adj_^2^=0.412, *p*=1.16×10^−26^), the combined FW+T1 model showed the most robust performance (R_adj_^2^=0.444, *p*=1.50×10^−29^). Similarly, all models significantly predicted longitudinal executive function slope (Actual: R_adj_^2^=0.424, *p*=9.20×10^−28^; FW: R_adj_^2^=0.404, *p*=5.89×10^−26^; T1: R_adj_^2^=0.420, *p*=2.38×10^−27^) and the combined FW+T1 model was the most robust (R_adj_^2^=0.446, *p*=1.13×10^−29^). When examining the age effect, we saw that actual age and all predicted brain ages each had a significant main effect on longitudinal memory slope (**Figure 6A**; Actual: β=−0.082, *p*=5.30×10^−10^; FW: β=−0.091, *p*=4.62×10^−11^; T1: β=−0.097, *p*=1.40×10^−8^), with the combined FW+T1 model showing the strongest relationship (β=−0.101, *p*=1.35×10^−11^). Likewise, we saw significant main effects for actual and all predicted brain ages on longitudinal executive function slope (**Figure 6B**; Actual: β=−0.128, *p*=1.58×10^−12^; FW: β=−0.125, *p*=1.20×10^−10^; T1: β=−0.163, *p*=4.25×10^−12^), with the combined FW+T1 model showing the strongest relationship (β=−0.158, *p*=1.65×10^−14^). We found no significant interactions between actual age or predicted brain ages and diagnostic status on longitudinal memory or executive function.

**Figure 6.**
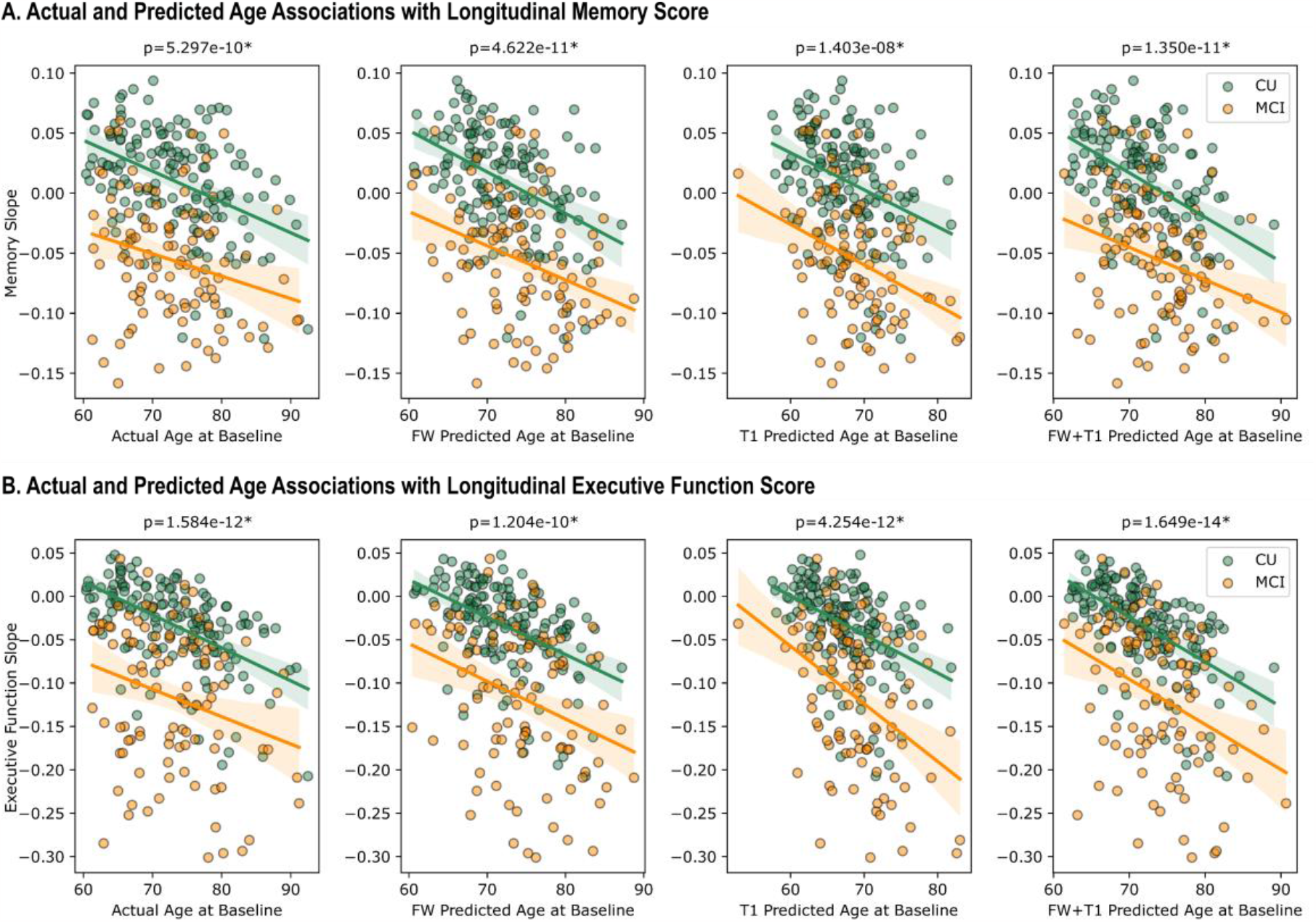
Actual and predicted age associations with longitudinal cognition. Actual and all derived ages are significantly associated with longitudinal memory (**A**) and executive function performance (**B**); FW+T1-derived age shows highest associations.

**Table 3** summarizes results of the competitive model analysis on longitudinal cognition. We found that covariates alone explained approximately 33% of the variance in longitudinal memory slope (R_adj_^2^=33.10%) and the addition of actual age led to an increase in overall model performance (ΔR^2^_adj_=9.68%). We then added predicted brain ages to this model one at a time to determine whether FW, T1, or FW+T1 predicted brain age contributed to any unique variance beyond covariates and actual age. We observed that all predicted brain ages were significant contributors to longitudinal memory slope and led to increases in R_adj_^2^ (FW: ΔR^2^_adj_=2.36%; T1: ΔR^2^_adj_=1.17%; FW+T1: ΔR^2^_adj_ =2.01%). Similarly, covariates alone explained approximately 30% of the variance in longitudinal executive function slope (R_adj_^2^=29.50%) and the addition of actual age led to a drastic increase in model performance (ΔR^2^_adj_ =12.99%). When iteratively adding each predicted brain age to the model to determine its unique contribution beyond covariates and actual age, we observed that all predicted brain ages explained additional variance in longitudinal executive function slope and led to increases in R_adj_^2^ (FW: ΔR^2^_adj_ =1.16%, T1: ΔR^2^_adj_ =2.54%; FW+T1: ΔR^2^_adj_ =2.67%).

**Table 3.** Comparison of Actual and Predicted Age Main Effects on Longitudinal Cognition.

|  | Longitudinal Memory Slope |  |  |  |  | Longitudinal Executive Function Slope |  |  |  |  |
| --- | --- | --- | --- | --- | --- | --- | --- | --- | --- | --- |
| | $\beta$ | SE | t | p | $\Delta R^2_{adj}$ | $\beta$ | SE | t | p | $\Delta R^2_{adj}$ |
| Covariates + Actual Age | -0.082 | 0.013 | -6.474 | <0.001 | 9.681 | -0.128 | 0.017 | -7.458 | <0.001 | 12.989 |
| Covariates + Actual Age + Predicted Age |  |  |  |  |  |  |  |  |  |  |
| FW | -0.060 | 0.018 | -3.372 | 0.001 | 2.362 | -0.060 | 0.025 | -2.441 | 0.015 | 1.160 |
| T1 | -0.057 | 0.016 | -3.558 | <0.001 | 1.174 | -0.097 | 0.028 | -3.482 | 0.001 | 2.539 |
| FW+T1 | -0.073 | 0.023 | -3.128 | 0.002 | 2.014 | -0.112 | 0.031 | -3.564 | <0.001 | 2.665 |

## 4. Discussion

The present study created 3 densely connected neural network models to predict brain age using FW, T1, and combined FW+T1 neuroimaging features, respectively. We evaluated model performance by comparing actual age with FW, T1, and FW+T1 predicted brain age then investigated the relationships between different age variables with cross-sectional and longitudinal cognitive performance (memory, executive function). Specifically, we examined age effects on baseline and longitudinal memory and executive function performance and conducted post hoc competitive model analyses to determine the unique contribution provided by each predicted brain age to variance in cognitive function. We report 3 main findings. First, we found that predicted brain ages from all 3 deep learning models using different sets of neuroimaging features (FW, T1, FW+T1) were highly associated with actual age; top neuroimaging features shown in model SHAP plots (**Figure 2**) were also biologically relevant to aging and cognitive decline, such as superior longitudinal fasciculus (SLF) FA_FWcorr_ and fornix FW in the FW model and thalamus and 3^rd^ ventricle in the T1 model. Second, we found that all predicted brain ages differentiated CU from MCI participants and significantly predicted both cross-sectional and longitudinal cognitive performance. Finally, we found that, among all 3 models, FW+T1 predicted brain age was the strongest predictor of cross-sectional and longitudinal cognitive performance and contributed the largest unique variance in these outcome variables.

### 4.1. Densely connected neural network robustly predicts age using neuroimaging features

We found that predicted brain ages generated by a densely connected neural network using 3 distinct sets of neuroimaging features (FW-corrected dMRI, T1-weighted MRI, combined FW+T1) all showed high correlation with actual age in baseline CU participants, which confirms findings from previous literature that have accurately predicted chronological age of healthy adults using neuroimaging-derived measures with machine learning approaches including deep learning^11,12,17,26–29^. Importantly, the top-contributing neuroimaging features identified for each model (**Figure 2**) provide biological interpretability as they include brain regions that have been associated with both normal aging and AD neuropathology. For instance, previous evidence has shown that thalamic volume, the most important feature identified in the T1 model, decreases with advancing age^30^ independently from total brain volume loss and correlates with cognitive speed and verbal memory performance^31,32^. Similarly, the identification of 3^rd^ ventricle volume as the second most important feature in the T1 model is consistent with prior literature which demonstrated that ventricular expansion is associated with normal aging and expands at an accelerated rate in individuals with cognitive impairment (MCI, AD)^33,34^ or AD-related pathology^35^. Among top features identified for the FW is the SLF, which is a white matter tract projecting from the occipital, parietal, and temporal lobes to the frontal cortex and is involved in language, attention, and memory^36^. Specifically, conventional FA within the SLF has been shown to undergo stable decline between ages 30-65 and accelerated decline after age 65^37^. Likewise, integrity of the fornix – a limbic white matter tract projecting from the hippocampus^38^– has been shown to decline with normal aging^39^ and to predict episodic memory^40^ and executive function performance^41^ in both healthy older adults and individuals with neurological disorders. Most existing literature on brain age prediction using machine learning techniques has leveraged T1-weighted MRI measures or conventional dMRI metrics. One significant advance in the present study is that we developed models using both T1-weighted and FW-corrected diffusion MRI data, and our results suggest that multi-modal MRI models may more accurately quantify brain age.

### 4.2. Predicted age is a more sensitive measure than actual age and predicative of cognition

We found that FW, T1, and FW+T1 predicted brain ages all differentiated CU from MCI patients by providing a significantly higher brain age for MCI patients even though the two groups did not differ in actual age, suggesting that predicted brain age may be a sensitive biomarker to AD clinical staging. This is consistent with previous research which computed predicted age difference (i.e., predicted brain age subtracted by chronological age) from T1-weighted MRI scans and found significantly larger predicted age difference in amnestic MCI participants compared with healthy controls^16^. Moreover, individuals with a higher predicted brain age at baseline were more likely to convert from MCI to AD^42^ or develop dementia later in life^18^. Studies generating predicted age difference from structural MRI scans of healthy controls have also found correlations with performance on traditional screening tools for AD (e.g., Mini-Mental State Examination, Clinical Dementia Ratio), anatomical measurements such as cortical thickness and hippocampal volume^43^, AD neuropathology such as β-amyloid positivity^16,26^, and AD risk factors such as *APOE*-ε4 carrier status^16,26^.

We also found that all predicted brain ages were robustly associated with cross-sectional and longitudinal cognitive function including baseline memory and executive function scores and longitudinal memory and executive function slopes. This agrees with prior literature that has found predicted age difference to be associated with memory and executive function impairment^16^ as well as early signs of cognitive decline^14^. However, the relationship between predicted age and both baseline and longitudinal cognitive function needs further clarification as one previous study found negative associations with psychomotor speed at baseline but no significant association with delayed recall performance or general cognitive status at baseline^44^. The present study supports predicted brain age as a sensitive biomarker along the AD continuum as it distinguishes between CU and MCI participants and is associated with memory and executive function performance at baseline and longitudinally.

### 4.3. Application of neural networks in clinical medicine

Deep learning algorithms, particularly neural networks, offer remarkable clinical utility by enabling researchers to harness complex patterns from large-scale data and consolidate this information into easy-to-use platforms. Prior neuroimaging studies have used deep learning methods to predict brain age^9,45–47^; however, the present study is the first to combine T1-weighted and FW-corrected diffusion MRI data, shedding light on the potential of using multi-modal MRI to accurately predict brain age and use it as an endophenotype for cognitive impairment and decline, especially in the context of aging and AD. Importantly, our neural networks add weight to the idea that both gray and white matter features are important to consider in aging and AD.

### 4.4. Strengths and limitations

The present study has several strengths, including a well-characterized longitudinal cohort with multi-modal MRI data with paired cognitive data. Regarding our neuroimaging analysis, one major strength is that we incorporated T1-weighted data in conjunction with FW-corrected diffusion MRI data, and this data was used as input into densely connected neural networks. Importantly, our data driven approach found that several aging related features (e.g., fornix integrity) were some of the highest contributing factors in our models. One limitation of this study is that it used a well-educated, mostly non-Hispanic white population, thus limiting our networks’ versatility. Future studies should incorporate more diverse populations to ensure that the neural networks are more generalizable. Moreover, although we have a large population with extensive longitudinal follow-up, one major limitation is that we only used data from a single cohort. Future studies leveraging multiple cohorts would drastically enhance our ability to predict brain age and likely improve its utility as an endophenotype for cognitive impairment and decline.

### 4.5. Conclusions

This study provided evidence that deep neural networks can be used to predict brain age, and that this predicted age is a strong predictor of cross-sectional cognitive impairment and future cognitive decline. Our findings provide evidence that using both T1-weighted and FW-corrected diffusion MRI data improves our ability to predict brain age; thus, future studies should consider both gray and white matter features when building deep learning models in aging and AD.

## 5. Acknowledgements

This study was supported by several funding sources, including K01-EB032898 (KGS), K01-AG073584 (DBA), U24-AG074855 (TJH), 75N95D22P00141 (TJH), R01-AG059716 (TJH), UL1-TR000445 and UL1-TR002243 (Vanderbilt Clinical Translational Science Award), S10-OD023680 (Vanderbilt’s High-Performance Computer Cluster for Biomedical Research). The research was supported in part by the Intramural Research Program of the National Institutes of Health, National Institute on Aging. Study data were obtained from the Vanderbilt Memory and Aging Project (VMAP). VMAP data were collected by Vanderbilt Memory and Alzheimer’s Center Investigators at Vanderbilt University Medical Center. This work was supported by NIA grants R01-AG034962 (PI: ALJ), R01-AG056534 (PI: ALJ), R01-AG062826 (PI: KAG), and Alzheimer’s Association IIRG-08-88733 (PI: ALJ).

